# Cyclic di-AMP regulates genome stability and drug resistance in Mycobacterium through RecA-dependent and -independent recombination

**DOI:** 10.1101/2024.05.13.593841

**Authors:** Sudhanshu Mudgal, Nisha Goyal, Kasi Manikandan, Rahul Saginela, Anusha Singhal, Soumyadeep Nandi, K. Muniyappa, Krishna Murari Sinha

## Abstract

In *Escherichia coli*, RecA plays a central role in the rescue of stalled replication forks, double-strand break (DSB) repair, homologous recombination (HR) and induction of the SOS response. While the RecA-dependent pathway is dominant, alternative HR pathways that function independently of RecA do exist, but relatively little is known about the underlying mechanism. Several studies have documented that a variety of proteins act either as positive or negative regulators of RecA to ensure high-fidelity HR and genomic stability. Along these lines, we previously demonstrated that the second messenger cyclic di-AMP binds to mycobacterial RecA proteins, but not *E. coli* RecA, and inhibits its DNA strand exchange activity *in vitro* via the disassembly of RecA nucleoprotein filaments. Herein, we demonstrate that *Mycobacterium smegmatis ΔdisA* cells, which lack c-di-AMP, exhibit increased DNA recombination, higher frequency of mutation and gene duplications during RecA-dependent and RecA-independent DSB repair. We also found that c-di-AMP regulates SOS response by inhibiting RecA-mediated self-cleavage of LexA repressor and its absence enhances drug resistance in *M. smegmatis ΔdisA* cells. Together, our results uncover a role of c-di-AMP in the maintenance of genomic stability through modulation of DSB repair in *M. smegmatis*.

**Significance:** Cyclic di-AMP is a second messenger present in bacteria and archaea and is implicated in a variety of functions in the cell, including DNA repair, cell wall metabolism, virulence, and gene expression. We show here that it maintains genome stability in Mycobacterium by regulating RecA-dependent and –independent DNA recombination pathways. It also regulates SOS response by inhibiting the self-cleavage of LexA by mycobacterial RecA. Absence of c-di-AMP leads to higher drug resistance in Mycobacterium.

## Introduction

Homologous recombination (HR) is an evolutionarily conserved process that plays a pivotal role in the maintenance of genome integrity in mitotic cells and homologous pairing and DNA strand exchange during meiosis (Gnügge and Symington, 2021, Bianco *et al*., 2001).In bacteria, HR is essential for DNA repair, reactivation of stalled replication forks, and acquisition of drug resistance (Kowalczykowski, 2015, Michel *et al*., 2007, Lusetti and Cox, 2002, Kuzminov and Stahl, 1999). Whereas HR in bacteria has been thought to be dependent on RecA-mediated HR pathways, a substantial level of HR occurs independently of RecA in certain genetic backgrounds (Bi *et al*., 1995, Dutra *et al*., 2007, Jain *et al*., 2021). In RecA-mediated HR, RecA protein preferentially binds to single-stranded DNA (ssDNA) to form a right-handed helical nucleoprotein filament, which subsequently promotes homologous pairing and DNA strand exchange (Flory *et al*., 1984, Kowalczykowski, 2015, Cox, 2007a).

The assembly of RecA nucleoprotein filament is rapid and bidirectional (Kowalczykowski, 2015, Cox, 2007b), although RecA polymerization is 50–60% faster in the 5′→3′ direction (Bell *et al*., 2012), and is tightly regulated by a network of positive (Muniyappa *et al*., 1984, Kowalczykowski and Krupp, 1987, Shan *et al*., 1997, Anderson and Kowalczykowski, 1997, Morimatsu and Kowalczykowski, 2003, Lusetti *et al*., 2004, Sakai and Cox, 2009) and negative (Venkatesh *et al*., 2002, Drees *et al*., 2004) protein effectors. Of note, mechanical forces regulate the stability of the RecA nucleoprotein filament and can counteract the inhibitory effect of RecX, prevent disassembly of the RecA filament, and stimulate the re-polymerization of RecA on ssDNA despite the presence of RecX (Le *et al*., 2014, Fu *et al*., 2013, Alekseev *et al*., 2022). We previously demonstrated that cyclic di-AMP (c-di-AMP) binds with low micromolar affinity to the C-terminus of mycobacterial RecA, but not to *E. coli* RecA, and inhibits its DNA strand exchange via disassembly of RecA nucleoprotein filaments (Manikandan *et al*., 2018).

The bacterial second messenger cdi-AMP, discovered in 2008 (Witte *et al*., 2008), is essential for the viability of various bacterial and archaeal species (Mudgal *et al*., 2021, Commichau *et al*., 2019, Braun *et al*., 2019). It has been implicated in diverse essential cellular processes including central metabolic pathways (Sureka *et al*., 2014, Krüger *et al*., 2022), osmolyte transport (Gundlach *et al*., 2019, Corrigan *et al*., 2013), cell wall homeostasis and drug resistance (Corrigan *et al*., 2011, Luo and Helmann, 2012), progression of sporulation (Oppenheimer-Shaanan *et al*., 2011), host–pathogen interactions and bacterial virulence (Woodward *et al*., 2010, McFarland *et al*., 2017) and DNA damage repair (Gándara and Alonso, 2015, Manikandan *et al*., 2018, Torres *et al*., 2019). In Mycobacteria, DisA is the sole diadenylate cyclase involved in the production of c-di-AMP and phosphodiesterase DhhP promotes its degradation (Manikandan *et al*., 2014, Yang *et al*., 2014). Little is known about the role of c-di-AMP in homology-directed DNA repair (HDR) pathways.

Following from our previous observation that c-di-AMP binds to the C-terminus of mycobacterial RecA and attenuates its DNA strand exchange activity *in vitro* through disassembly of RecA nucleoprotein filaments (Manikandan *et al*., 2018), we demonstrate in this study that cells lacking c-di-AMP exhibit: (i) significantly increased levels of HDR and inappropriate HR, (ii) high frequency of deletions and duplications and (iii) higher level of SOS induction and drug resistance. Our results highlight a previously unappreciated, crucial role for c-di-AMP in the regulation of HR, acquisition of antibiotic resistance, and maintenance of genome stability in *M. smegmatis*.

## Results

### Absence of c-di-AMP potentiates HDR-mediated DSB repair in *M. smegmatis*

In general, cells utilize two major mechanisms to repair DSBs, namely NHEJ and HR. In addition, DSBs are also repaired by two other alternative pathways: single-strand annealing (SSA) and the alt-NHEJ/microhomology-mediated repair (Scully *et al*., 2019). To gain insight into the relative contributions of NHEJ and HR pathways to DSB repair in mycobacteria, the Shuman and Glickman group had developed a reporter system in which I-SceI-induced DSB repair could be studied genetically and mapped physically (Gupta *et al*., 2011). To elucidate the physiological relevance of c-di-AMP during DSB repair in *M. smegmatis*, we leveraged the same I-SceI-based reporter assay (Fig. 1). In this genetic assay, repair of DSB yields blue and white colonies: white colonies can result from NHEJ-mediated repair and LacZ^+^ (blue) colonies from error-free HR or SSA repair pathway (Fig. 1). Our results revealed that the frequency of LacZ^+^ colonies in the mutant *ΔdisA* strain was approximately three times higher as compared to the WT strain: this implies that increased HR-mediated DSB repair is likely due to the absence of c-di-AMP (Fig. 2A).These results are also consistent with the results of our previous study where we had shown that c-di-AMP attenuates the DNA strand exchange activity of mycobacterial RecA (Manikandan *et al*., 2018). *ΔdisA M. smegmatis* cells had undetectable level of RecA initially but the expression came back to the WT level with time and has been used in the present study (Fig. S3 C, D, E).

**Fig. 1.**
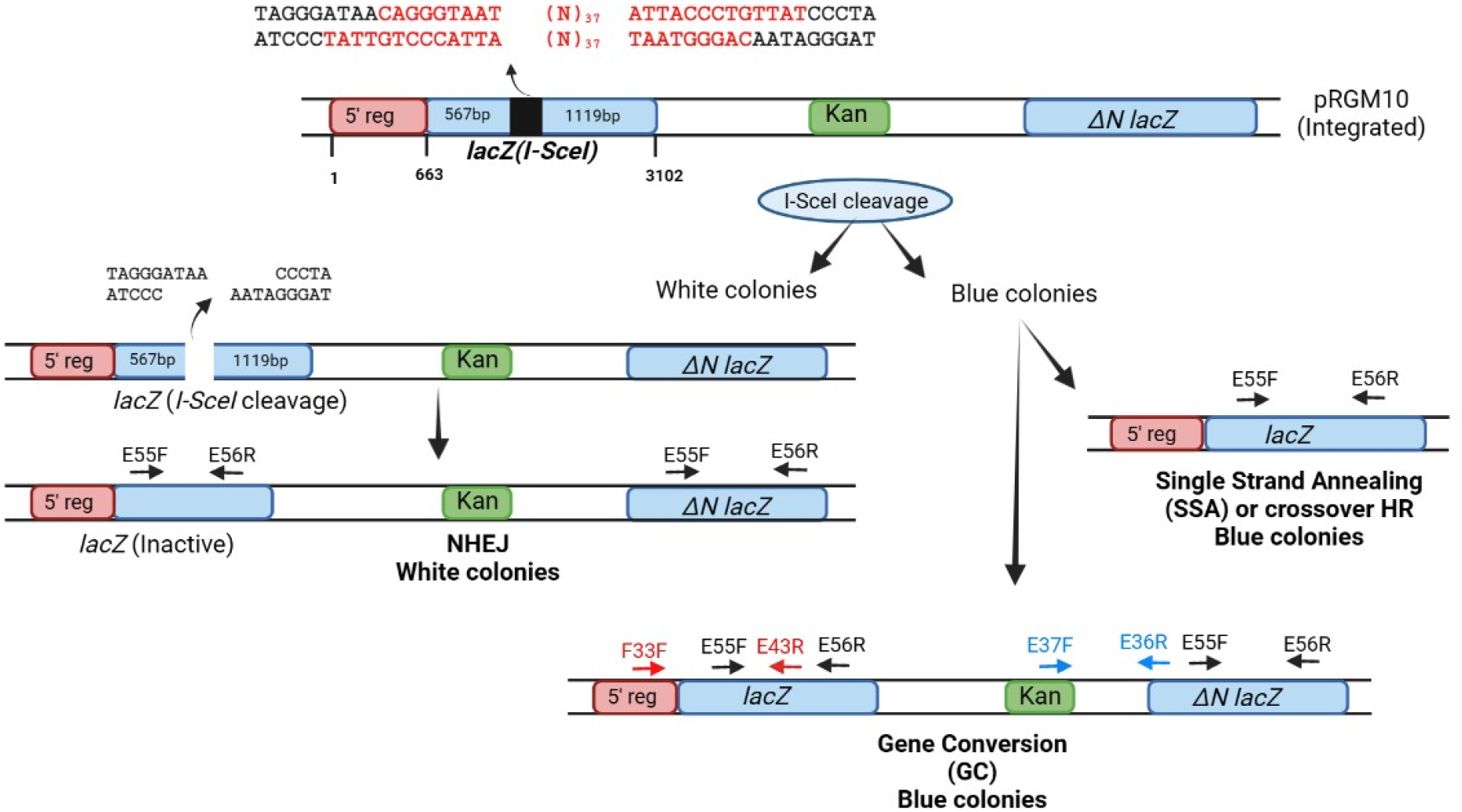
Reporter construct pRGM10 and the known repair pathways and outcome: The upper panel shows the reporter construct pRGM10 integrated at the *attB* site of *M. smegmatis* cells. The sequence written above pRGM10 are the two 18-mer I-SceI target sequences placed 37 nt apart in opposite orientation in *lacZ*(*I-SceI*). The sequence shown in black are of the DSB generated with 3’-overhangs after cleavage by I-SceI. The break will be repaired either by NHEJ or HR giving rise to scorable white and blue colonies respectively. The repair outcomes were analyzed by doing PCR amplification with diagnostic primers. The position and orientation of the primers with their numbers have been shown with arrows to indicate their orientation on the repaired (outcome) sequence.PCR with primers E55F and E56R will give a product of 1311 bp on repair by GC or SSA (blue colonies) whereas repair by NHEJ will give two PCR products of 1311 bp and 779 bp or smaller as per the number of nucleotides deleted. Primer F33F was used to detect the presence of the 5’ *lacZ* DNA sequence. PCR with F33F and E43R was used to detect repair outcome shown in Fig. 3B. E37F and E36R were used to detect kanamycin gene (Kan) to distinguish between GC and SSA.

**Fig. 2.**
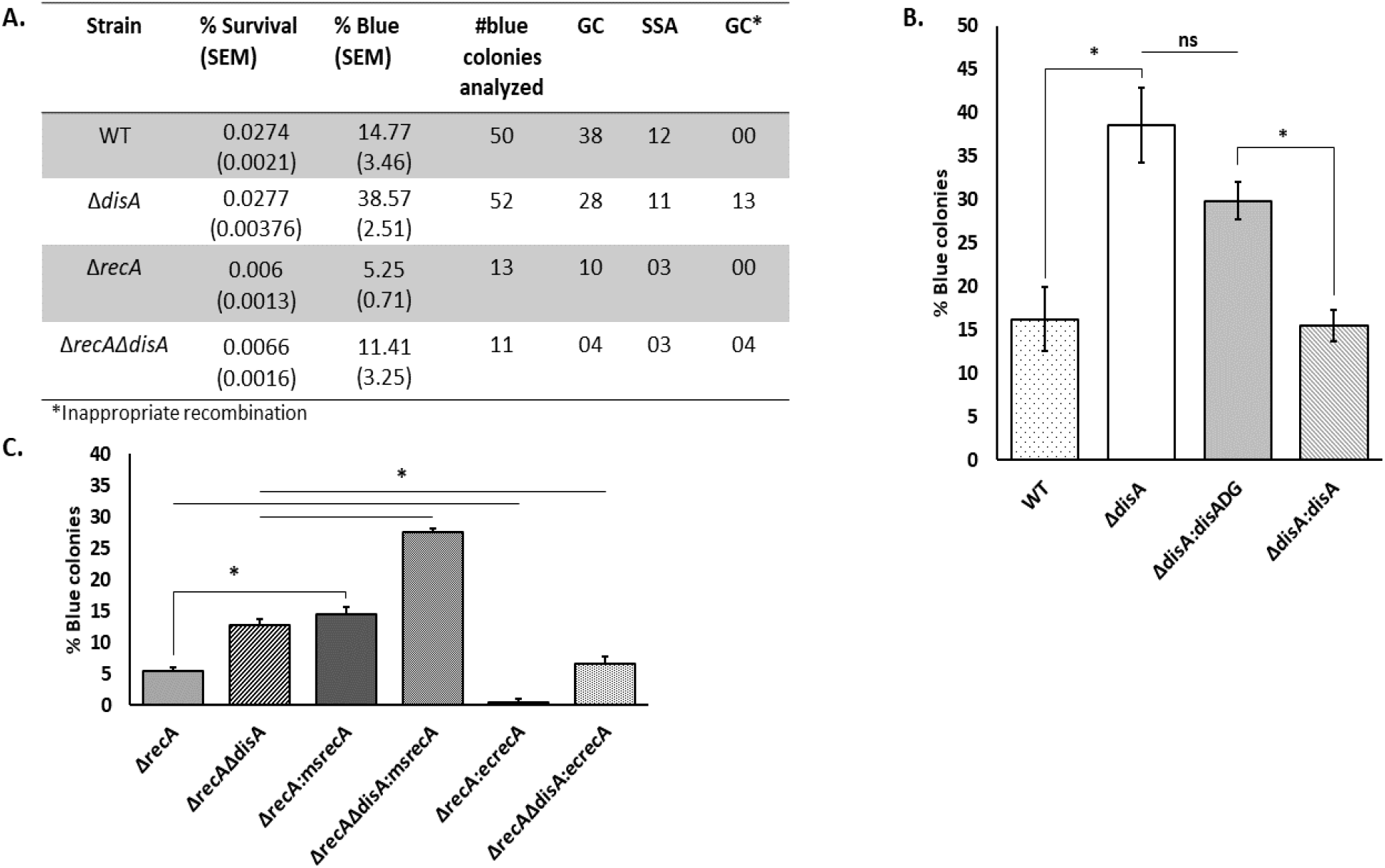
Absence of c-di-AMP leads to higher homologous recombination: A) The table shows the % survival and % blue colonies (with SEM) of the different mycobacterial strains after repair of I-SceI-induced DSB. The number of blue colonies analyzed to determine the repair outcome and the repair by GC, SSA, and GC^*^ as determined has been given. GC^*^ represents the blue colonies obtained after the repair of DSB by NHEJ as explained in the results. B) Bar graph showing the percentage of blue colonies obtained in WT, *ΔdisA, ΔdisA:disADG (ΔdisA* complemented with active site mutant DisA) and *ΔdisA:disA (ΔdisA* complemented with wild type DisA). The percentage of blue colonies was calculated as: number of blue colonies/total survivors on the plate x 100. The difference among the blue colonies in different strains was found to be statistically significant (\**p*≤0.05). C) Percentage of blue colonies obtained in WT, *ΔrecA, ΔrecAΔdisA, ΔrecA:msrecA* (*ΔrecA* complemented with MsRecA), *ΔrecAΔdisA:msrecA* (*ΔrecAΔdisA* complemented with MsRecA), *ΔrecA:ecrecA* (*ΔrecA* complemented with EcRecA) and *ΔrecAΔdisA:ecrecA* (*ΔrecAΔdisA* complemented with EcRecA). Each experiment was done with at least three biological replicates and each replicate was performed in triplicate.

To test whether the increased HR-mediated DSB repair was due to the lack of c-di-AMP and not DisA protein, *ΔdisA* cells were complemented with either WT *disA* or catalytically inactive *disA*^D72A/G73A^ variant (designated as DisADG) (Manikandan *et al*., 2014) and the repair efficiency was determined by counting LacZ^+^ colonies. We observed that the percentage of LacZ^+^ colonies in the *ΔdisA* and *disADG* strains increased approximately 2- to 3-fold as compared to the WT (Fig. 2B). On the other hand, the percentage of LacZ^+^ colonies in *ΔdisA* cells bearing WT *disA* allele was similar to that of wild-type cells. Collectively, these results support the idea that c-di-AMP negatively affects HR-mediated DSB repair *in M. smegmatis*.

### Cyclic di-AMP regulates both the RecA-dependent and -independent HR

Given that the deletion of *disA* led to an increase in the amount of LacZ^+^ colonies, we asked whether the increase in LacZ^+^ colonies is because of higher DNA strand exchange activity of RecA in the absence of c-di-AMP as predicted from our previous work (Manikandan *et al*., 2018). For this purpose, *M. smegmatis ΔrecA* and *ΔrecAΔdisA* knockout mutants were constructed and their ability to form LacZ^+^ colonies was compared with that of *ΔdisA* cells. Consistent with our premise, a 3.4 fold decrease in the number of blue LacZ^+^ colonies was observed in the double mutant *ΔrecAΔdisA* mutant strain compared to the *ΔdisA* cells, suggesting that RecA-mediated HR pathway is involved in the formation of blue LacZ^+^ colonies in *ΔdisA* cells (Fig. 2B, 2C). It is interesting to note that blue colonies were obtained even in the absence of RecA in *ΔrecAΔdisA* cells suggesting the presence of RecA-independent HDR (Lovett *et al*., 2002, Dutra *et al*., 2007, Jain *et al*., 2021). This result was also confirmed in *ΔrecA* cells where blue colonies were obtained though the percentage decreased significantly (3 fold) compared to the WT *M. smegmatis* cells (Fig. 2A, 2C). Interestingly, the percentage of blue colonies in *ΔrecAΔdisA* double mutant is more than 2 fold higher than in *ΔrecA* cells (Fig. 2C) suggesting that c-di-AMP also regulates RecA-independent homology mediated DSB repair.

Complementation of *ΔrecA* and *ΔrecAΔdisA* cells with cognate *M. smegmatis* RecA (MsRecA) increased the HDR to WT and *ΔdisA* levels respectively but complementation with *E. coli* RecA (EcRecA) did not increase the percentage of HDR. The extent of homology mediated repair in EcRecA complemented cells was as less as the parent cells (Fig. 2C) suggesting that EcRecA is not functional in *M. smegmatis*.

### Deletion of *disA* does not affect NHEJ repair

Next, white colonies were sequenced to decipher the role of c-di-AMP on DSB repair promoted by NHEJ. The DSB repair junctions were amplified with forward and reverse primers, E55F and E56R (Fig. 1) and PCR products were sequenced to detect the NHEJ-mediated repair outcome. Of the thirty representative white-colored colonies sequenced for the WT and *ΔdisA* strain each, 52% were repaired through NHEJ and 48% had intact I-SceI site suggesting inactivation of the *I-SceI* gene through mutation (Gupta *et al*., 2011). Both the I-SceI target sequences were cleaved in all the NHEJ colonies and most of the repair was done by deleting nucleotides at the break site to expose homology at the ends followed by ligation. The deletions were mostly of a few nucleotides and they were similar in both the WT and *ΔdisA* cells (Supplementary fig. S1A, B). The surprising outcome was the absence of NHEJ in *ΔrecAΔdisA* cells. We sequenced 20 colonies and 19 were found to have I-SceI inactivation and 1 colony had an insertion of 5 nucleotides at the 3’ end (Supplementary fig. S1C). So, there is a possibility that c-di-AMP/DisA might have indirect effect on NHEJ or impact NHEJ mediated repair under certain conditions.

### Deletion of *disA* results in hyper-recombinogenic RecA protein

The function of RecA is tightly regulated by a complex network of positive and negative modulators that act to maintain a balance between inappropriate and appropriate recombination events (Venkatesh *et al*., 2002, Bakhlanova *et al*., 2016, Kowalczykowski, 2015, Cox, 2007a). Since c-di-AMP attenuates mycobacterial RecA promoted DNA pairing and strand exchange reactions (Manikandan *et al*., 2018), we reasoned that RecA may display a characteristic “hyper-rec” phenotype with aberrant recombination events in *ΔdisA* cells. To test this hypothesis, the repair outcome across I-SceI junction in *lacZ(I-SceI)* gene in blue colonies were PCR amplified with the primers F33F and E43R (Fig. 3B). In all the cases, a major PCR product migrating at the expected size of 1507 bp was observed; this results from the DSB repair via GC or SSA (Gupta *et al*., 2011). Unexpectedly, however, an additional band of about 780-bp was seen in 25% of blue LacZ^+^ colonies (Fig.3A). Sequencing showed that the 780-bp PCR product originated from NHEJ-repaired *lacZ(I-SceI)* gene (Fig.3C). Since the DSB repair by NHEJ should produce white colonies, the 1507-bp PCR product was also sequenced to decipher the possible mechanism behind the formation of blue LacZ^+^ colonies (Fig. 3D). The results revealed that the 662-bp region located near the 5’ end of *lacZ(I-SceI)* gene has recombined with the downstream (*ΔNlacZ*) gene giving rise to blue LacZ^+^ colonies in this assay. Although this recombination event is similar to the repair of DSBs by GC, we posit that the DSB was first repaired by NHEJ, followed by HR by ‘hyper-rec’ RecA, which differs from the previous observations (Gupta *et al*., 2011).

**Fig. 3.**
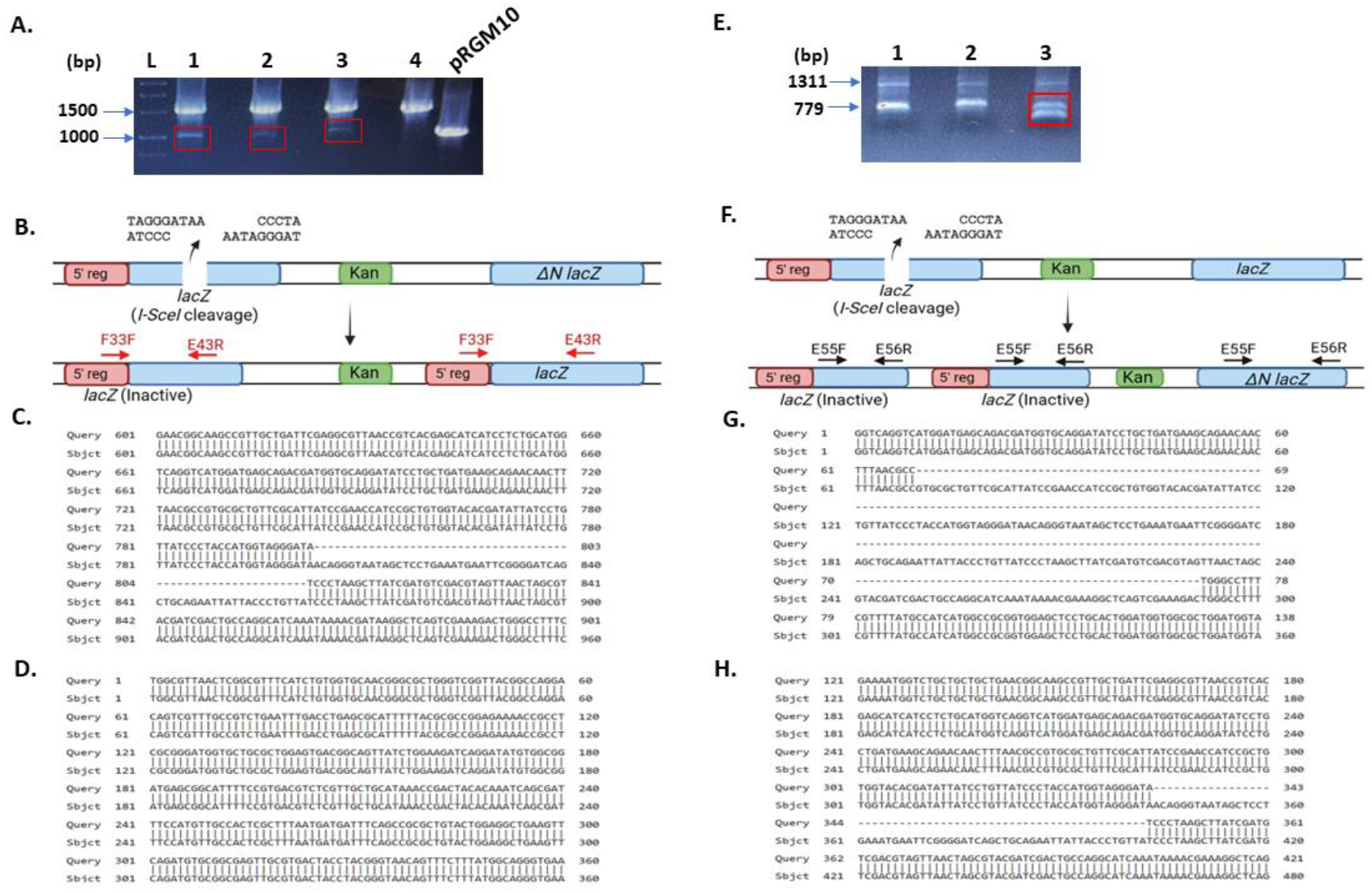
Hyper-recombination and gene duplication in *ΔdisA* cells: A. Repair events of 4 representative blue colonies (numbered 1-4) have been shown. #1-3 have undergone repair by GC^*^ whereas #4 is repaired by GC after I-SceI cleavage. The colonies were analyzed by PCR amplification with primers F33F and E43R shown in panel B. PCR products were separated on an agarose gel and ethidium bromide-stained gel has been shown. A major product of 1507 bp was obtained along with an additional smaller product (shown in red box). pRGM10 was used as a control template. Arrows on the left side of the gel show the size and position of the bands. B. Figure showing the repair outcome of the blue colonies #1-3. The break has been repaired by NHEJ but GC has also occurred transferring the 5’ 662nt to *ΔNlacZ* giving rise to active *lacZ*. C. The smaller product results from nucleotide deletion across the break site during NHEJ. The figure shows alignment of the sequence of smaller PCR product of colony 1 with that of the *lacZ-ISceI* sequence. The gaps in the alignment arise because of nucleotide deletion during NHEJ repair of DSB. D. Sequence of the bigger PCR product (with primers F33F and E43R) confirms the recombination of 5’ 662nt to *ΔNlacZ*. E. Repair events in 3 representative white colonies after I-SceI expression were analyzed by PCR amplification using primers E55F and E56R. Two products of 1311 and around 779 bp were obtained as expected (as explained in Fig. 1) but there was an additional smaller product in colony #3. Both the smaller bands of colony #3 have been marked with red box. Arrows on the left indicate the size and position of the bands. F. Figure showing the repair outcome in colony # 3. There is duplication of the *lacZ(I-SceI)* gene after repair by NHEJ. G and H. The sequence alignment of the smaller PCR products with the *lacZ* gene has been shown. The gaps indicate deletion of different lengths.

We also interrogated whether such NHEJ-repaired blue colonies were obtained in *ΔrecA* and *ΔrecAΔdisA* strains. To this end, 13 and 11 blue colonies from strains with a single *ΔrecA* mutation and *ΔrecAΔdisA* double mutations, respectively, were sequenced. Of these, 36% of blue LacZ^+^ colonies in *ΔrecAΔdisA* strain were repaired by NHEJ followed by HR between the 5’ end of *lacZ(I-SceI)* and *ΔNlacZ*, but no such recombination events occurred in the *ΔrecA* or WT strains. This result along with the results in the previous section clearly suggests that a significant percentage of HR events can proceed independently of RecA in *M. smegmatis* and that c-di-AMP inhibits both the RecA-dependent and RecA-independent HR, although the precise mechanism remains unclear.

The outcome of DSB repair in white colonies was also assessed by PCR amplification of their genomic DNA using primers E55F and E56R (Fig. 1A). As expected, two PCR products migrating at the expected size of 1311 bp and 779 bp or smaller depending on the deletion during NHEJ were observed. In one of the 30 white colonies analyzed from the *ΔdisA* strain, we observed the 1311 bp product but got 2 smaller products instead of one (Fig. 3E). Sequencing of the smaller products suggested that they were generated because of NHEJ deletion (Fig. 3G, H). We reasoned that the second product would appear either because of duplication of the first repaired gene or a mixed colony. To rule out the second possibility, we streaked the culture of the colony and determined the repair outcome in 6 individual colonies by doing genomic PCR as above (Supplementary Fig.S2A) All the six colonies gave 2 smaller products as above suggesting that the colony was not mixed but there was a duplication of *lacZ(I-SceI)* gene.

### Deletion of *disA* enhances the SOS response and promotes drug resistance

Most bacteria utilize the tightly regulated RecA/LexA SOS response system to induce DNA repair genes, wherein activated RecA stimulates the autocleavage of LexA repressor (Foster, 2007). Mycobacteria harbor at least two independent DNA damage response (DDR) pathways: the LexA/RecA-independent SOS response (Movahedzadeh *et al*., 1997, Durbach *et al*., 1997, Davis *et al*., 2002, Rand *et al*., 2003) and the PafBC-regulated pathway (Müller *et al*., 2018, Adefisayo *et al*., 2021, Brzostek *et al*., 2021). We have demonstrated previously that c-di-AMP binds to the C-terminus of mycobacterial RecA and inhibits DNA strand exchange through disassembly of RecA nucleoprotein filaments (Manikandan *et al*., 2018).Hence, we postulated that *ΔdisA* cells might possess hyper-recombinogenic RecA, which may cause strong SOS induction and mutations, resulting in drug resistance. To this end, the SOS response was assessed in the *M. smegmatis ΔdisA* mutant after UV irradiation and growth on rifampicin (Boshoff *et al*., 2003). We found a 2-fold increase in the number of rifampicin-resistant colonies in the *ΔdisA* mutant as compared to the WT, whereas rifampicin-resistant colonies were undetectable in *ΔrecA* and *ΔrecAΔdisA* knockout mutants (Fig. 4A): this implies that the hyper-rec activity of RecA induces a hypermutable state that promotes acquisition of resistance to rifampicin (Boshoff *et al*., 2003, Adefisayo *et al*., 2021). Furthermore, complementation of *M. smegmatis ΔrecA* mutant with its cognate WT *recA*, but not *E. coli recA*, conferred rifampicin-resistance. Similarly, *ΔrecAΔdisA* knockout mutant, complemented with its cognate WT *recA* formed higher number of rifampicin-resistant colonies, indicating a role for c-di-AMP in the SOS response and mutagenesis (Fig. 4A). In contrast, *ΔrecA* and *ΔrecAΔdisA* mutants complemented with *E. coli recA* did not induce the SOS response (Fig. 4A), providing evidence for a direct link between c-di-AMP and suppression of RecA-induced mutagenesis.

**Fig. 4.**
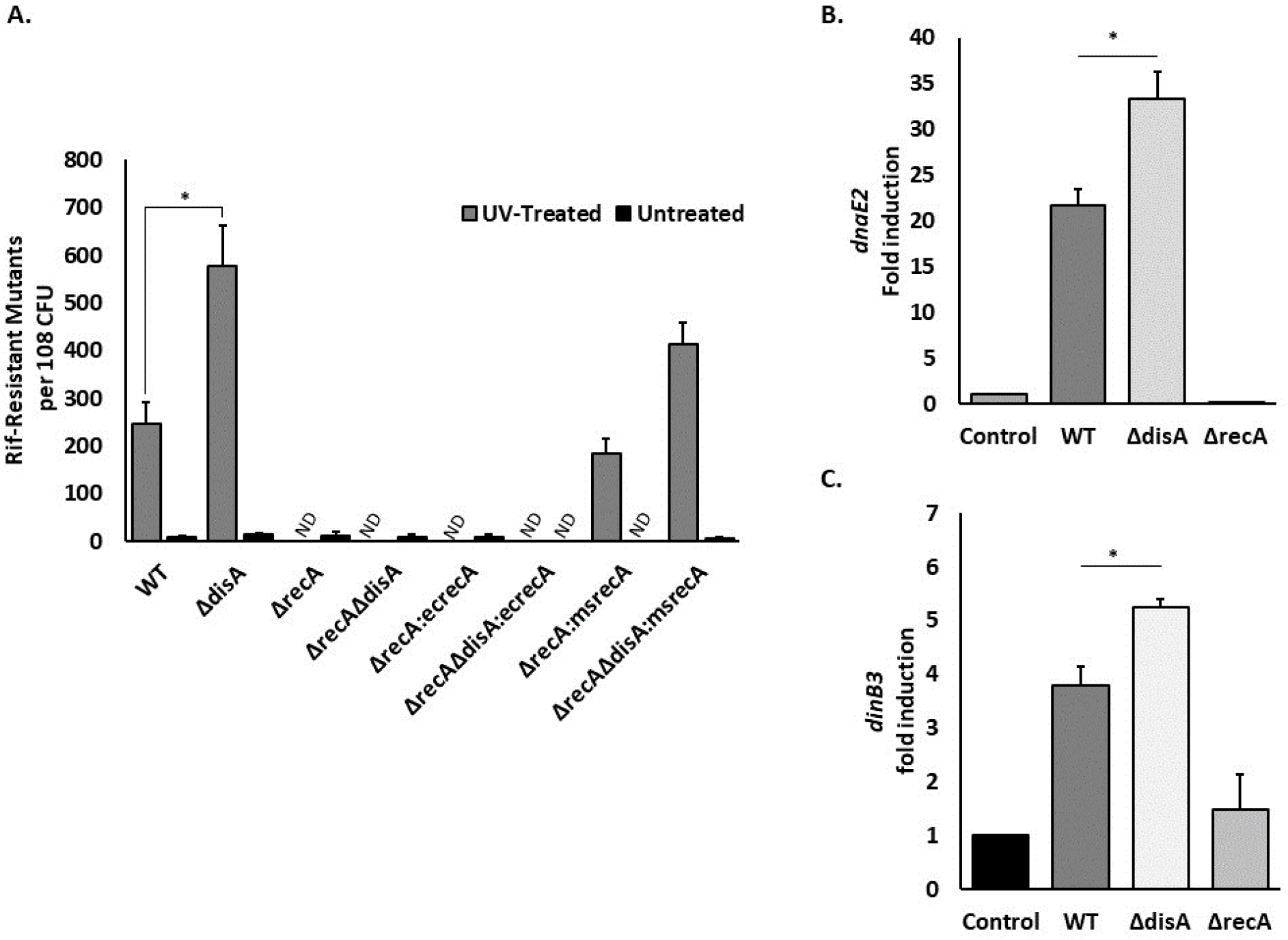
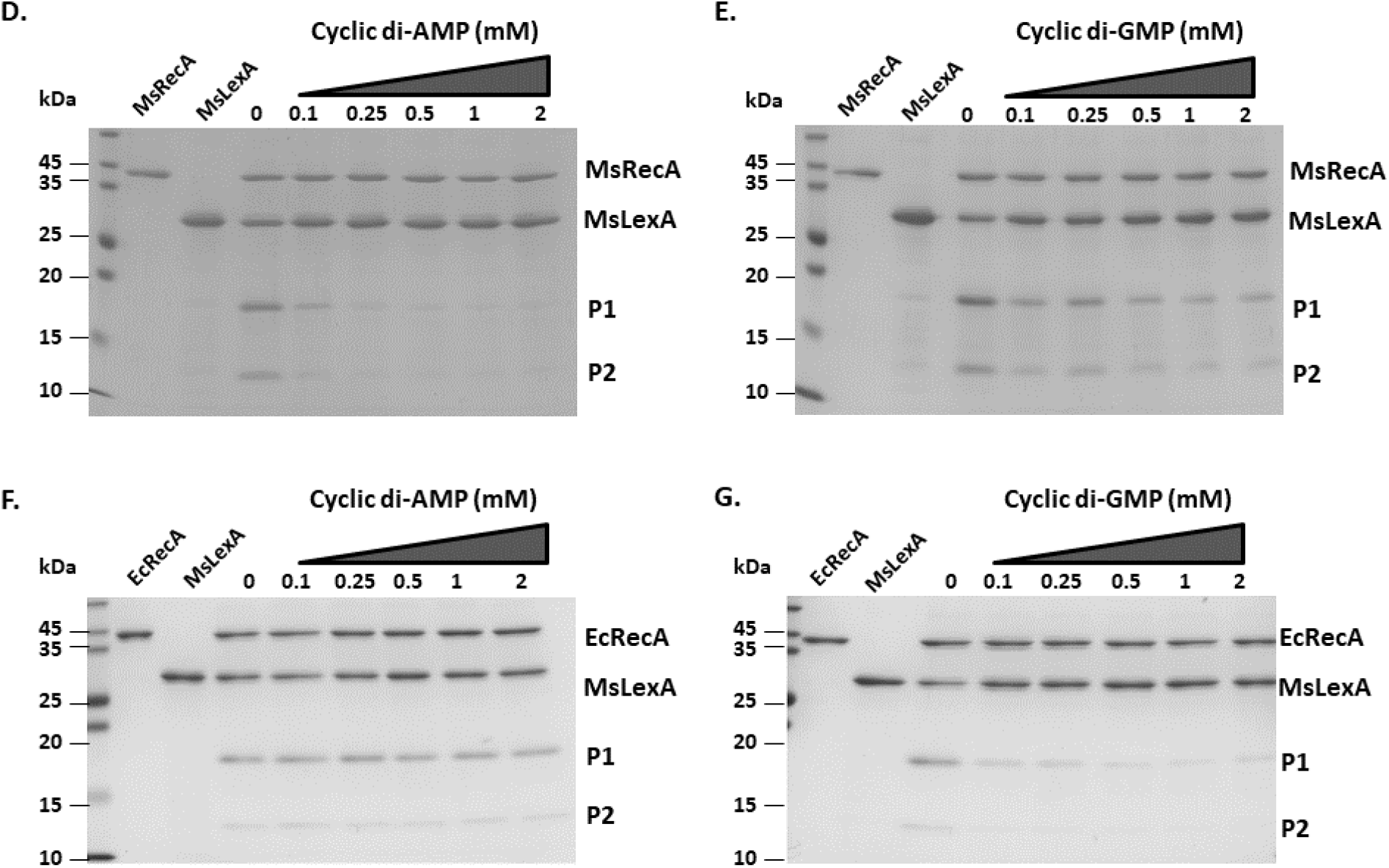
Absence of cyclic di-AMP induces stronger SOS response. A.Frequency of rifampicin-resistant mutants per 10^8^ cells in untreated (black bars) or UV-treated (grey bars) of the indicated *M. smegmatis* strains. ND, not detected. **B and C**. Transcript levels of *dnaE2* and *dinB3* in WT, *ΔdisA*, and *ΔrecA M. smegmatis* cells after treatment with quinolones to induce SOS response. **\***p < 0.05 for indicated comparisons by paired t-test. Error bars represent SD. All experiments were done with 3 biological replicates and each replicate was done in triplicate. **D-G: Inhibition of LexA cleavage by c-di-AMP and c-di-GMP:** Fig.D and E show the cleavage of MsLexA by MsRecA in the presence of c-di-AMP and c-di-GMP respectively. MsLexA was incubated with MsRecA in the presence of increasing concentration of c-di-AMP (C) and c-di-GMP (D) as described in the methods. The reaction mixture was run on SDS-PAGE and the coomassie-stained gel has been shown. The first two lanes show the purified MsRecA and MsLexA recombinant proteins as labeled at the top. The different concentration of the dinucleotides used in the reaction varies from 0 to 2 mM as mentioned at the top of the lanes. The position and size of protein ladder has been marked at the left. P1 and P2 (marked on the right) are the two cleavage products of MsLexA. Fig. F and G show the cleavage MsLexA with EcRecA in the presence of c-di-AMP and c-di-GMP respectively as shown at the top of the figures. The cleavage assay was carried out as above and coomassie-stained SDS-PAGE gel has been shown.

In mycobacteria, trans-lesion synthesis (TLS) DNA polymerases, DnaE2 and DinB, enable cells to confer rifampicin resistance (Boshoff *et al*., 2003, Dupuy *et al*., 2022). During SOS response *dnaE2* is induced in a RecA-dependent manner, although the cell intrinsic/extrinsic factor involved in the induction of *dinB* remain unclear. To gain insights into the mechanism underlying acquisition of drug resistance by *ΔdisA* mutant, we monitored the induction of *dnaE2, dinB1, dinB2* and *dinB3* in response to 10 µg/ml of ofloxacin treatment, followed by qPCR. The results show a 20- and 35-fold increase in the transcript levels of *dnaE2* in WT and *ΔdisA* mutant, respectively (Fig. 4B). On the other hand, a 3.7- and 5.3-fold increase in the *dinB3* transcript levels were observed in the WT and *ΔdisA* cells respectively.

Next, we sought to characterize the biochemical basis of SOS-induced mutagenesis promoted by hyper-recombinogenic activity of *M. smegmatis* RecA (MsRecA) in the *ΔdisA* mutant. To this end, MsRecA-mediated autocleavage of its cognate LexA repressor was monitored in the absence and presence of c-di-AMP. The reaction products were analysed by SDS-PAGE and visualized by staining with Coomassie blue. It revealed that c-di-AMP, but not c-di-GMP, abrogates MsRecA-mediated autocleavage of LexA repressor in a concentration-dependent manner (Fig. 4C-D). In contrast, c-di-GMP inhibited EcRecA-mediated autocleavage of *M. smegmatis* LexA repressor but c-di-AMP did not (Fig. 4E-F).

## Discussion and Perspectives

DNA integrity scanning protein or DisA was discovered as a protein that scans the chromosome during sporulation in *B. subtilis* and blocks sporulation on detecting any damage (Bejerano-Sagie *et al*., 2006). The DAC activity of DisA is inhibited on binding to Holliday junction or replication fork like structures (Witte *et al*., 2008). *B. subtilis* and Mycobacterium contain a conserved operon consisting of *disA* and *radA* genes suggesting a role of DisA/c-di-AMP in DNA damage repair. Several lines of evidence suggest that c-di-AMP plays a crucial role in different cellular processes such as antibiotic tolerance, biofilm formation and pathogenesis (Commichau et al., 2015). Despite these findings, key questions remain unanswered: for example, what are the target effectors that are responsible for such diverse effects of c-di-AMP and the underlying mechanisms by which it exerts its effects. To this end, our previous work has demonstrated that c-di-AMP binds to the C-terminus of mycobacterial RecA, but not EcRecA, and attenuates DNA strand exchange activity of MsRecA *in vitro* (Manikandan *et al*., 2018). Here we show that *M. smegmatis ΔdisA* strain, which lacks c-di-AMP, exhibits increased HDR levels of RecA-dependent and RecA-independent DSB repair, although the precise role of c-di-AMP in latter pathway remains unclear, warranting further investigation. Of note, *ΔdisA* null mutant displays hyper-recombination and hypermutable phenotype, which are important mechanisms responsible for acquired drug resistance in bacteria. Further analysis of *ΔdisA* mutant revealed that it exhibits higher frequency of deletions and duplications and a 2-fold increase in the acquisition of rifampicin resistance. Moreover, we found that the loss of c-di-AMP results in robust SOS response via inhibition of RecA-mediated self-cleavage of LexA repressor. The inhibition of LexA cleavage in both Mycobacterium and *E. coli* suggests that the regulation of SOS response by cyclic dinucleotides could be a general phenomenon. Together, our results uncover a role of c-di-AMP in the maintenance of genomic stability through modulation of DSB repair in *M. smegmatis*.

Although considerable progress has been made towards understanding of the role of accessory proteins that negatively regulate RecA-mediated HR (Venkatesh *et al*., 2002, Lusetti *et al*., 2004; Cox 2007), our knowledge about the potential roles of endogenous small molecule inhibitors, including bacterial second messenger(s), in the regulation of RecA function is limited. In this context, it is worth highlighting that c-di-AMP, but not c-di-GMP, inhibits the recombination-like activities of mycobacterial RecA proteins, but not the EcRecA, *in vitro* (Manikandan *et al*., 2018), consistent with the *in vivo* data reported in this study. The combined *in vitro* and *in vivo* data provide valuable insights into the molecular basis by which c-di-AMP regulates RecA activities and DNA repair/recombination functions. Taking into consideration the effects of c-di-AMP *in vivo* and *in vitro*, we propose a model as to how it may regulate genomic stability in *M. smegmatis* (Figure 5). Given the presence of cyclic dinucleotides in all domains of life, their role in maintaining genome stability might be conserved. There are suggestions of the involvement of cGAS in genome stability in metazoans (Chen *et al*., 2020).

**Fig. 5.**
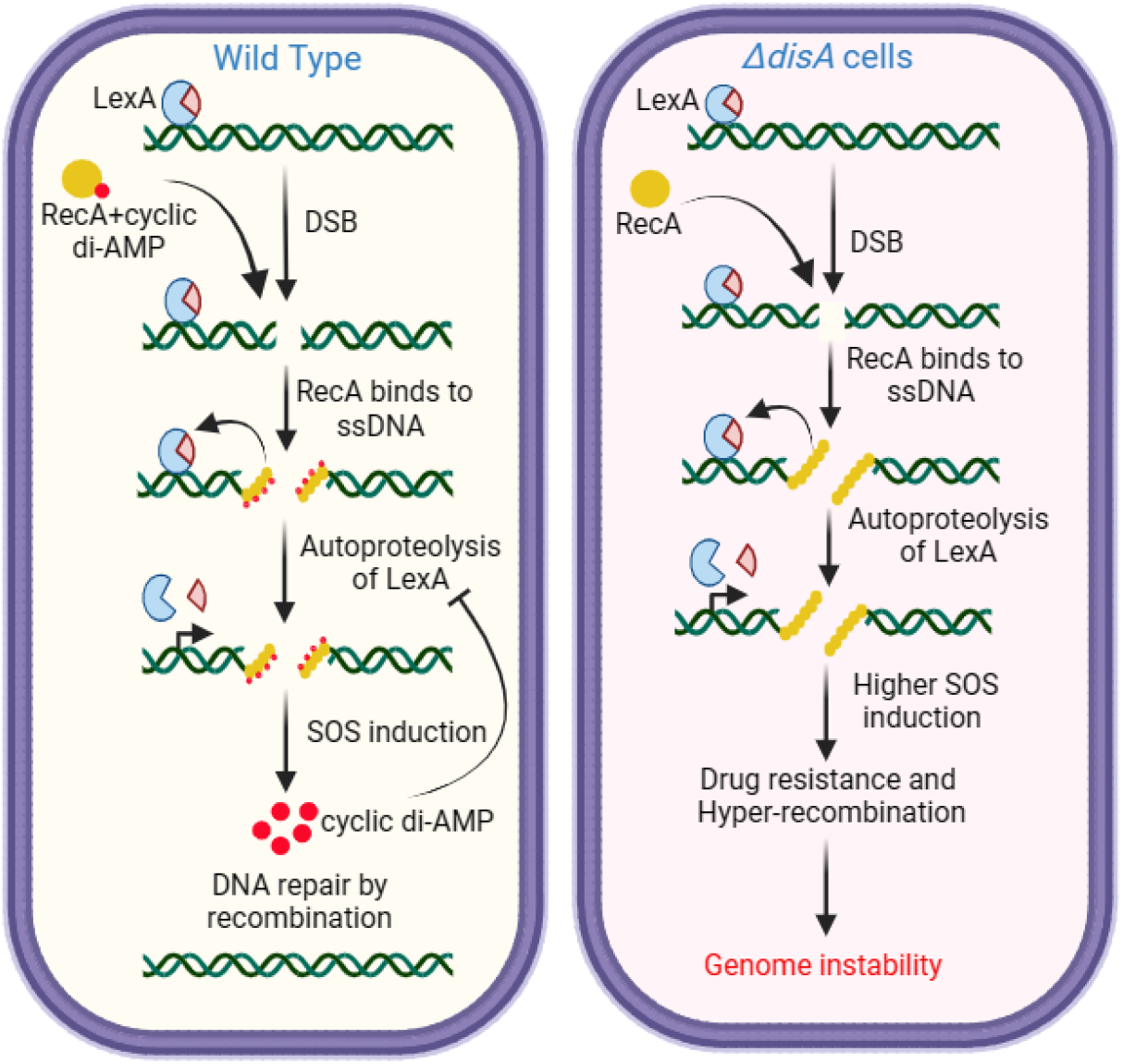
Model showing the function of c-di-AMP in RecA-dependent HDR and SOS response. C-di-AMP inhibits the formation of RecA nucleoprotein filament leading to subdued SOS induction whereas there is a higher amount of recA nucleoprotein filament and SOS induction in the absence of c-di-AMP. *disA* transcript is induced as a part of SOS response, c-di-AMP so synthesized in turn regulates SOS response, homologous recombination, and genome stability. In the absence of c-di-AMP, there is higher SOS induction, drug resistance, unnecessary recombination, gene duplication, and genome instability.

Bacterial resistance to antibiotics is a major global health concern, which poses a serious and rapidly growing threat to human, animal, and environment health (Salam *et al*., 2023). The enzymes involved in DNA repair/recombination have been linked to the emergence of drug-resistant pathogenic bacteria, especially multidrug-resistant and extensively drug-resistant TB strains (Gygli *et al*., 2017). Consequently, attenuation of the SOS response system via preferential inhibition of RecA has been proposed as a possible therapeutic target to block the development of multi-drug resistance and resistance-conferring mutagenesis (Mo *et al*., 2016, Thi *et al*., 2011). Several inhibitors of RecA activities *in vitro* have been developed including ATP-competitive nucleotide analogs (Lee *et al*., 2005,Wigle *et al*., 2006), polysulfated naphthyl compounds (Wigle and Singleton, 2007, Nautiyal *et al*., 2014, Zhou *et al*., 2021), as well as short peptides (Yakimov *et al*., 2017). To our knowledge, none of these synthetic small molecules have demonstrated RecA-specific biological activities in bacterial cell cultures. This work provides a framework for designing or re-engineering endogenous small molecules with enhanced activity against bacterial RecA to reduce the acquisition of antibiotic resistance mutations and evolution of drug-resistant bacteria.

## Materials and methods

Reporter construct pRGM10 and I-SceI mediated DSB: Reporter construct pRGM10 was used to study I-SceI mediated DSB repair in this study (Gupta *et al*., 2011). The construct was integrated at the *attB* site of *M. smegmatis* chromosome by electroporating pRGM10 along with pBSIntegrase which contains the L5 integrase gene (Manikandan *et al*., 2018). Cells were plated in the presence of 20 µg/ml of kanamycin on 7H10 agar plates supplemented with 0.5% glycerol and 10% OADC. Integration was confirmed by doing PCR with suitable primers. The forward primer (D69F) anneals 5’ to the *attB* site in the chromosome whereas the reverse primer E35R anneals 5’ to *lacZ(I-SceI)* gene within the construct.

ORF of *I-SceI* was amplified from pMSG375 using the primers E16F and E17R and cloned between SphI-EcoRI site of pVV17 to give the plasmid pVVI-SceI. To induce DSB, *M. smegmatis* cells containing pRGM10 were electroporated with pVVI-SceI and the colonies were selected against (100 µg/ml) hygromycin on 7H10 agar plates containing 40 µg/ml of X-gal. Outcome of the repair was determined by doing genomic PCR with suitable primers across the junction and sequencing the PCR product. The genomic DNA of blue colonies obtained because of the repair by GC or SSA were PCR amplified with forward primer that anneals within the first 662 nt of *lacZ* ORF (F33F) and the reverse primer E43R anneals downstream of the break site (Fig. 1). This set of primers will give a single PCR product of 1507 bp. The repair outcome of GC and SSA were distinguished by doing PCR with kanamycin specific primers E37F and E36R which will give a PCR product of 1442 bp for repair by GC.

The repair outcomes of white colonies were determined by sequencing the PCR product amplified by diagnostic primers that anneal across the junction. The colonies with intact I-SceI site were sequenced to confirm mutations in *I-SceI* gene by amplifying the gene with suitable primers.

### Complementation of the mutant cells

The complementation constructs in this study have been made in the plasmid pYS2 and integrated at the intergenic region of *msmeg_5848* and *msmeg_5849* (Gupta *et al*., 2011) through DNA recombineering as described before (Shenkerman *et al*., 2014). The construct contains 5’ and 3’ homologous sequences for integration and a cassette containing the gene to be expressed is placed in between the homologous sequences (Supplementary Fig.S3) The 5’ homologous region containing the last 721 nt of *msmeg_5848* was amplified from *M. smegmatis* genome using the primers F23F and F24R, digested with SpeI and SwaI and cloned in similarly digested pYS2. 3’ homologous region containing the terminal 726 bp of *msmeg_5949* was similarly PCR amplified with primers F58F and F59R, digested with NsiI and PacI and cloned in similarly digested plasmid construct containing the 5’ homologous region.

The genes to be expressed were first placed under hsp promoter by cloning them initially in pVVgfp(5) (Manikandan *et al*., 2018). The gene cassettes were then amplified from these constructs with the forward primer F19F which anneals at the beginning of the hsp promoter and a gene specific reverse primer which anneals over the termination of the ORF. The reverse primers used to amplify *msrecA, ecrecA* and *ecmsrecA* are F22R, F21R and F22R respectively. Both the forward and the reverse primers contain SwaI site. The PCR product thus obtained contains the hsp promoter along with the gene ORF, digested with SwaI and cloned at the SwaI site of the construct containing the 5’ and 3’ homologous regions. The construct places the gene 5’ to the *gfp-hyg* cassette which is flanked by loxP on either side. The plasmid was digested with SpeI and NsiI to release the complementing construct. The construct was electroporated in *M. smegmatis* cells (either WT or different mutants) which expresses Che9c 60-61 from the plasmid pYS1. The electroporated cells were plated on 7H10 containing hygromycin and sucrose and grown at 42°C to cure them of pYS1. The cells so obtained have the complementing construct integrated at the intergenic region of *msmeg_5848* and *msmeg_5849* and were screened based on the expression of GFP. The integration and orientation of the gene was further confirmed by doing PCR with primers F48F and F49R (Fig. S3B). The complemented cells were then transformed with pML2714 which expresses P1 Cre recombinase to remove *gfp-hyg* cassette. pML2714 was subsequently removed by growing the cells at 42°C. The expression of the genes was confirmed by western blotting (Fig. S3C-E).

### UV-induced mutagenesis

For UV-induced mutagenesis, cells were grown in 7H9 medium until saturation. The cultures were then diluted to OD600 = 0.02 and grown further to OD600 = 0.5. 5.0 mL of the cultures were transferred to 90 x 15 mm sterile plastic petri plates, and exposed to 20 mJ/cm^2^ UV radiation using a UVP CL 1000 Ultraviolet Crosslinker. The treated cells were transferred to 5 mL of fresh 7H9 media and grown in a shaker incubator at 37 °C with shaking at 150 RPM for 3 hours. The untreated control of each strain was processed similarly. 100 µL of cells from each sample were plated in duplicate on 7H10 agar plates containing 10% OADC, 0.5% glycerol, and 80 µg/ml of rifampicin, and plates were wrapped in a foil to prevent potential effects of photolyase, incubated at 37°C for 72 hours to determine rifampicin-resistant CFU. Cells were taken from each sample and appropriate dilutions were plated on 7H10 agar plates containing 10% OADC and 0.5% glycerol in the absence of antibiotic to determine viable CFU. The resistant mutants obtained on rifampicin plates were normalized to viable CFU. Each set of experiments was done with 3 biological replicates; each replicate was done in triplicate.

Change in the transcript level of DnaE2 and DinB polymerases during SOS induction was assessed by doing RT-qPCR. *M. smegmatis* cells were grown to OD600≈0.5 and treated with 10 µg/ml of ofloxacin for 3 hours. The cells were harvested by centrifugation and the cell pellet was re-suspended in RNAprotect bacteria reagent (Qiagen) and incubated at RT for 5 min. The cells were re-centrifuged and the pellet was used for total RNA extraction using RNeasy kit (Qiagen) as per the manufacturer’s protocol. Briefly, the pellet was suspended in buffer RLT. The cells were sonicated, centrifuged and the lysate was collected. 1 volume of 70% ethanol was mixed with the lysate, and the mixture was promptly transferred onto the RNeasy mini spin column. The column was then centrifugated for 15s at 8000*g*, and the flow-through was discarded. Subsequently, the column was washed with buffer RW1 and buffer RPE. Finally, RNA was eluted from the column with RNase-free water. The eluted RNA (1 μg) was treated with DNase I and subjected to cDNA synthesis using RevertAid H minus reverse transcriptase (Thermo Scientific) with random hexamer. cDNA prepared was used for RT-qPCR reactions which was performed using TaqMan Assay. *ΔΔ*CT values of each strain were calculated using 16S rRNA as control.

### LexA cleavage assay

LexA cleavage assay was performed as per the published protocol (Patil *et al*., 2011, Wipperman *et al*., 2018). The reaction was carried out in a 10 µl reaction mixture. 1 µM MsRecA was first incubated with shown amount of c-di-AMP for 5 minutes at 37°C followed by the addition of 20 mM Tris-Cl, pH 7.0, 8 mM MgCl2, 1 mM DTT, 3 mM ATPϒS and 5 µM SS DNA (63-mer oligonucleotide, HJ03). The reaction mixture was further incubated at 37°C for 5 minutes followed by the addition of 3 µM LexA and the reaction was carried out for 5 minutes at 37°C. The cleavage was assessed by running the reaction mixtures on SDS-PAGE and coomassie staining.

## Data availability statement

The data that support the findings presented in the manuscript have been included in the manuscript and supplementary information. Any additional data can be requested from the corresponding author.

## Acknowledgement

The authors thank Prof. Michael Glickman of Memorial Sloan Kettering Cancer Center for providing reporter constructs pRGM9 and pRGM10, Prof. Eyal Gur of Ben-Gurion University of the Negev for the gene deletion plasmids pYS1, pYS2, pML2714, Dr. Anjana Badrinarayanan of National Centre for Biological Sciences for E. coli RecA antibodies and Prof. Stewart Shuman of Memorial Sloan Kettering Cancer Center for his inputs on the manuscript. The authors are thankful to Drs. Ranjita Ghosh Moulick and Kaustav Bandopadhyay of Amity University Haryana for access to their equipments. Work in KMS laboratory is supported by funding from SERB (CRG/2020/003946). SM is an ICMR SRF (TB-fellowship/14/2022-ECD-I) and Kasi Manikandan is UGC-DSKPDF.

## Notes

### Competing Interest Statement

The authors have declared no competing interest.

