## Supplementary Information for "Cyclic di-AMP regulates genome stability and drug resistance in Mycobacterium through RecA-dependent and -independent recombination"

**Plasmids used in this study**:

| Name | Description | Source |
| --- | --- | --- |
| pRGM10 | Reporter construct, Plasmid carrying two inactive *lacZ* genes, [*lacZ(ISce-I)* and *∆NlacZ*], two ISce-I sites in opposite orientation, *attP* site, Km^R^ | (Gupta *et al.*, 2011) |
| pYS1 | A shuttle vector with a tm^S^ mycobacterial origin of replication, Che9c 60-61 protein, counter-selectable marker SacB, acetamide inducible promoter, Km^R^ | (Shenkerman *et al.*, 2014) |
| pYS2 | Plamsid carrying two *loxP* sites flanking the *gfp-hyg* cassette, Amp^R^(for *E. coli*), Hyg^R^ | (Shenkerman *et al.*, 2014) |
| pML2714 | Plasmid carrying tm^S^ mycobacterial origin of replication, express P1 Cre recombinase, Km^R^ | (Shenkerman *et al.*, 2014) |
| pdisAGFP | Plasmid carrying *M. tb disA* (*rv3586*) cloned in pVVGFP at HindIII and KpnI site, Km*^R^*, expresses DisA with C-terminal GFP tag | This work |
| pdisAGFP(2) | DisA mutated at D72A and G73A in pdisAGFP | This work |
| pRecA(1)gfp | Plasmid carrying *M. smegmatis recA* (*msmeg_2723*) cloned in pVVGFP at HindIII and KpnI site | (Manikandan *et al.*, 2018) |
| pEcRecA(1)gfp | Plasmid carrying *E. coli recA* cloned in pVVGFP at HindIII and KpnI site | (Manikandan *et al.*, 2018) |
| pVV17 | NdeI site in pVV16 mutated to SphI, Hyg^R^ | This work |
| pVVI-SceI | SphI-EcoRI digested *I-SceI* gene from pMSG375 cloned at similarly digested pVV17, Hyg^R^ | This work |
| pBSIntegrase | L5 integrase gene cloned at XhoI/EcoRI site of pBS (pBlueScript) plasmid | (Manikandan *et al.*, 2018) |
| pYS2MsRecA | Complementation construct for *M. smegmatis recA* (*msmeg_2723*), *msmeg_2723* along with hsp promoter amplified from pRecA(1)gfp and cloned in pYS2. | This work |
| pYS2EcRecA | Complementation construct for *E. coli recA* (*ecrecA*), *ecrecA* along with hsp promoter amplified from pEcRecA(1)gfp and cloned in pYS2. | This work |
| pYS2disA | Complementation construct for *M. tb disA*(*mtbdisA*), *mtbdisA* along with hsp promoter amplified from pdisAGFP and cloned in pYS2. | This work |
| pYS2disA(2) | Complementation construct for *M. tb disA* D72AG73A mutant in pYS2. | This work |

Km^R^= kanamycin resistant, Amp^R^=Ampicillin resistant, Hyg^R^= Hygromycin resistant, tm^S^= Temperature sensitive, *gfp-hyg*= Green fluorescent protein and hygromycin resistant, SacB= Sucrose, DG= D72AG73A.

**Bacterial strains used in this study:**

| Name | Description | Source |
| --- | --- | --- |
| *∆disA* | Derivative of *M. smegmatis* mc^2^ 155 carrying an unmarked deletion in *disA* | (Manikandan *et al.*, 2018) |
| *∆recA* | Derivative of *M. smegmatis* mc^2^ 155 carrying an unmarked deletion in *recA* | (Manikandan *et al.*, 2018) |
| *∆recA∆disA* | Derivative of *M. smegmatis* mc^2^ 155 carrying unmarked deletions in *disA* and *recA* | This work |
| mc^2^155:pRGM10 | mc^2^ 155 derivative carrying Km resistance plasmid vector pRGM10 integrated at the *attB* locus | This work |
| *∆disA*:pRGM10 | *∆disA* derivative carrying plasmid vector pRGM10 integrated at the *attB* locus | This work |
| *∆recA*:pRGM10 | ∆*recA* derivative carrying plasmid vector pRGM10 integrated at the *attB* locus | This work |
| *∆recA∆disA*:pRGM10 | *∆recA∆disA* derivative carrying plasmid vector pRGM10 integrated at the *attB* locus | This work |
| *∆recA*:*msrecA*:pRGM10 | Derivative ∆*recA* complemented with *M. smegmatis recA (msrecA*) at the intergenic region of *msmeg_5848* and *msmeg_5849* and plasmid pRGM10 integrated at *attB* locus | This work |
| *∆recA∆disA*:*msrecA*:pRGM10 | Derivative of *∆recA∆disA* complemented with *M. smegmatis recA (msrecA*) at the intergenic region of *msmeg_5848* and *msmeg_5849* and plasmid pRGM10 integrated at *attB* locus | This work |
| *∆recA*:*ecrecA*:pRGM10 | Derivative of ∆*recA* complemented with *E. coli recA* (*ecrecA)* at the intergenic region of *msmeg_5848* and *msmeg_5849* and plasmid pRGM10 integrated at *attB* locus | This work |
| *∆recA∆disA*:*ecrecA*:pRGM10 | Derivative of *∆recA∆disA* complemented with *E. coli recA* (*ecrecA)* at the intergenic region of *msmeg_5848* and *msmeg_5849* and plasmid pRGM10 integrated at *attB* locus | This work |
| *∆disA*:*mtbdisA*:pRGM10 | Derivative of *∆disA* complemented with *M. tb disA* (*mtbdisA)* at the intergenic region of *msmeg_5848* and *msmeg_5849* and plasmid pRGM10 integrated at *attB* locus | This work |
| *∆disA*:*mtbdisADG*:pRGM10 | Derivative of *∆disA* complemented with *M. tb disADG* (*mtbdisADG)* at the intergenic region of Msmeg_5848 and Msmeg_5849 and plasmid pRGM10 integrated at *attB* locus | This work |

**List of primers used in this study:**

| Name | Primer Sequence (5’ to 3’) | Description |
| --- | --- | --- |
| E16F  E17R | TTTTTTGCATGCAGAAAGGAGGCCATATGGG  TTTTGAATTC TTATTTCAGG AAAGTTTCGG AG | To amplify *I-SceI* from pMSG375 and cloning in pVV16 |
| E27F  E28R | GGAATCACTTCCATATGCCCAAGACAATTGCGGATC  GATCCGCAATTGTCTTGGGCATATGGAAGTGATTCC | To mutate NdeI and create an SphI site in pVV16. To mutate G388 to C in pVV16 and create pVV17a |
| E29F  E30R | CCGGAGGAATCACTTCCGCATGCCCAAGACAATTGC  GCAATTGTCT TGGGCATGCG GAAGTGATTC CTCCGG | To mutate AT385 to GC in pVV17a to create pVV17 |
| E37F  E36R | CACCGGATTCAGTCGTCACTC  CTCCAGTACAGCGCGGCTG | Anneals downstream of *kan* and N-terminal of lacZ of pRGM10. |
| E42F  E43R | GCTGATTCGAGGCGTTAACCG  TTCACCGCTTGCCAGCGGC | To confirm deletion after DSB repair. Binds across the break site in *lacZ(I-SceI)* |
| E55F  E56R | TGCCGATCGCGTCACACTAC  CAGACGCCACTGCTGCCAG | To confirm deletion after DSB repair. Binds across the break site in *lacZ(I-SceI)* |
| F19F  F21R | TTTTTTATTTAAATGGTGACCACAACGACGCGC  TTTTTTATTTAAATTTAAAAATCTTCGTTAGTTTCTGCTACG | To amplify *ecrecA* with hsp promoter from pEcrecA(1)gfp |
| F22R | TTTTTTATTTAAATTCAGAAGTCAACCGGGGCCGG | To amplify *msrecA* along with hsp promoter from pRecA(1)gfp |
| F23F  F24R | TTTTACTAGTCCCGTGCGGATCTTCCCC  TTTTTTATTTAAATCTAGGGTGCGTCGGTCAG | Primers to amplify 3' *msmeg_5848* |
| F33F | GAAAGCTGGCTACAGGAAGGC | Primer anneals downstream of 5’-region of pRGM10 |
| F48F  F49R | CTTCTCGCGCGTCGTCGCG  TGTCCTCGGCCCTCCGATC | To confirm the complementation, F48F anneals 721 bp upstream of the termination of msmeg_5848, F49R anneals at 107-125 nt of the hsp promoter |
| F58F  F59R | TTTTTAATTAACTAGGCCTGCAGCTTCTCGAAC  TTTTATGCATGATCAGGTGGGGCTGCGCG | Primers to amplify 3' *msmeg_5849* |

F and R at the end of primer number indicate forward and reverse primer respectively**.**

**TAGGGATAACAGGGTAAT(N)_37_ATTACCCTGTTATCCCTA**

**ATCCCTATTGTCCCATTA(N)_37_TAATGGGACAATAGGGAT**

A.) **Wild-Type**

```
TACCATGGTAGGGATA  Δ1/Δ1     TCCCTAAGCTTATC
ATGGTACCATCCCT            ATAGGGATTCGAATAG
```

```
TACCATGGTAGGGATAAT  /Δ2    CCCTAAGCTTATC
ATGGTACCATCCCTAT         TAGGGATTCGAATAG
```

```
GATGGTGC       Δ115/Δ4     CCCTAAGCTTATC
CTACCACG                   GGGATTCGAATAG
```

B.) ***∆disA***

```
TACCATGGTAGGGATA  Δ1/Δ1   TCCCTAAGCTTATC
ATGGTACCATCCCT          ATAGGGATTCGAATAG
```

```
TACCATGGTAGGGAT  Δ2/    TATCCCTAAGCTTATC
ATGGTACCATCCC         TAATAGGGATTCGAATAG
```

**TACCATGGTAGGGATAACA	/Δ6	CTAAGCTTATC	ATGGTACCATCCCTATTGT	GATTCGAATAG**

```
CATTATCC            Δ57/Δ5   CTAAGCTTATC
GTAATAGG                     GATTCGAATAG
```

```
TATCGTGCGGTGGT  Δ297/Δ219   AACTGCCTGAAC
ATAGCACGCCACCA              TTGACGGACTTG
```

C.) ***∆recA∆disA***

```
CATGGTAGGGATAACAGGG     TATCCCTAAGCTT
GTACCATCCCTAT          AATAGGGATTCGAA
```

**Fig. S1. Molecular outcome of DSB reapir by NHEJ:** The sequence at top are the two 18-mer I-SceI target sequences placed 37 nt apart in opposite orientation in *lacZ*(*I-SceI*). The sequence shown in red are of the DSB generated with 3’ non-complementary overhangs after cleavage by I-SceI. ∆n/∆n represents the number of nucleotides deleted from the 3’ overhangs during repair, inserted nucleotides have been shown in green. A.) and B.) shows the repair outcomes in wild-type and *∆disA* cells. C.) shows the insertion of nucleotides at the break site, there is no deletion of nucleotides during repair.

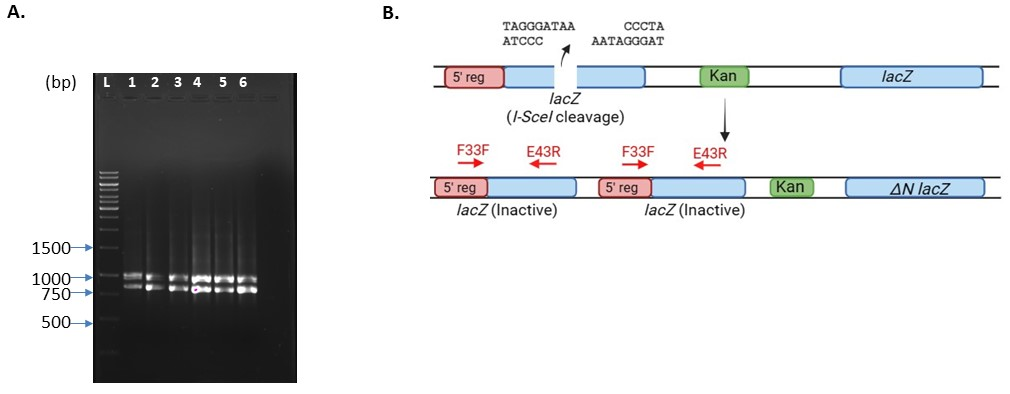

**Fig. S2 Duplication of *lacZ(I-SceI)* gene:**

**A)** PCR product of six different white colonies obtained after streaking. Genomic PCR amplification of 6 white colonies was done with primers F33F and E43R. PCR product was run on agarose gel and visualized by ethidium bromide staining as shown. **B)** The upper panel shows the I-SceI cleaved reporter construct pRGM10 whereas the lower figure shows its repair outcome. Duplication of *lacZ(I-SceI)* gene giving rise to 2 inactive *lacZ* during repair has been shown.

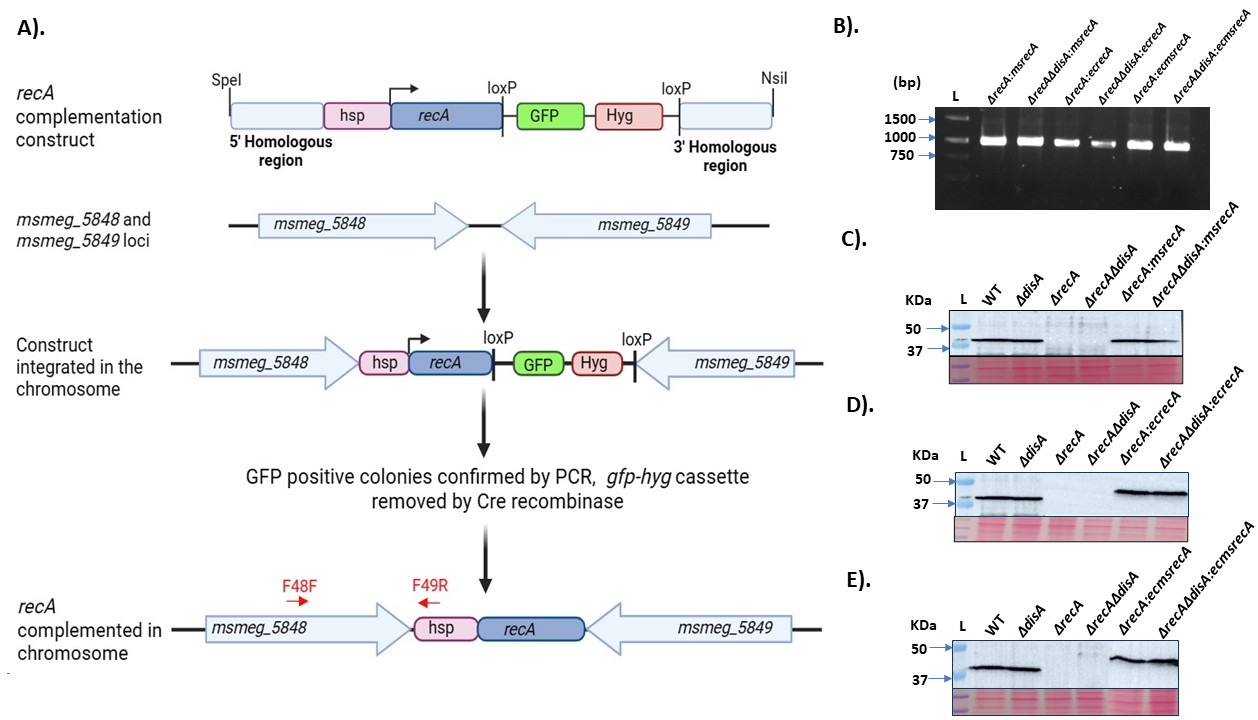

**Fig. S3 Complementation of mutants: A.** Construct for the complementation of *recA/disA* in *M. smegmatis*. Genes which are to be complemented have been placed under the heat shock promoter (hsp) for expression. The recombineering construct was integrated at the intergenic region of *msmeg_5848* and *msmeg_5849* in the genome. The *gfp-hyg* cassette was later removed by electroporating a plasmid expressing Cre recombinase. Integration of the construct was confirmed by doing PCR with primers F48F and F49R. The arrows show the orientation and position of the primers. **B.** Ethidium bromide-stained agarose gel showing the genomic PCR products of different strains as mentioned above the respective lanes. **C. D and E.** Western blot showing the expression of RecA. Cell extract (soluble) of the indicated strains were separated on 12% SDS-PAGE and probed with *anti-*RecA antibodies. The blot shown in C was probed with anti-MsRecA. D and E were probed with commercially available *E. coli* *anti*-RecA (Abcam 6379). The ponceauS-stained membrane shows equal loading of the extract.
